# Modulations of local synchrony over time lead to resting-state functional connectivity in a parsimonious large-scale brain model

**DOI:** 10.1101/2021.01.20.427443

**Authors:** Oscar Portoles, Yuzhen Qin, Jonathan Hadida, Mark Woolrich, Ming Cao, Marieke van Vugt

## Abstract

Biophysical models of large-scale brain activity are a fundamental tool for understanding the mechanisms underlying the patterns observed with neuroimaging. These models combine a macroscopic description of the within- and between-ensemble dynamics of neurons within a single architecture. A challenge for these models is accounting for modulations of within-ensemble synchrony over time. Such modulations in local synchrony are fundamental for modeling behavioral tasks and resting-state activity. Another challenge comes from the difficulty in parametrizing large scale brain models which hinders researching principles related with between-ensembles differences. Here we derive a parsimonious large scale brain model that can describe fluctuations of local synchrony. Crucially, we do not reduce within-ensemble dynamics to macroscopic variables first, instead we consider within and between-ensemble interactions similarly while preserving their physiological differences. The dynamics of within-ensemble synchrony can be tuned with a parameter which manipulates local connectivity strength. We simulated resting-state static and time-resolved functional connectivity of alpha band envelopes in models with identical and dissimilar local connectivities. We show that functional connectivity emerges when there are high fluctuations of local and global synchrony simultaneously (i.e. metastable dynamics). We also show that for most ensembles, leaning towards local asynchrony or synchrony correlates with the functional connectivity with other ensembles, with the exception of some regions belonging to the default-mode network.

**Author summary:** Here we present and evaluate a parsimonious model of large-scale brain activity. The model represents the brain as a network-of-networks structure. The sub-networks describe the neural activity within a brain region, and the global network encodes interactions between brain regions. Unlike other models, it capture progressive changes of local synchrony and local dynamics can be tuned with one parameter. Therefore the model could be used not only to model resting-state, but also behavioural tasks. Furthermore, we describe a simple framework that can deal with the arduous task of identifying global and local parameters.

## Introduction

The human brain is one of the most complex systems found in nature, consisting of billions of neurons. Human behavior cannot be understood from only the computing properties of individual neurons. Instead, human behavior requires the coordination of many ensembles of neurons at multiple spatial scales. Neuroimaging studies reveal that patterns of large-scale local and distributed coordination appear and dissolve during behavioral tasks [1–5], as well as resting-state [6–10].

Biophysical models of large-scale brain networks provide a unified analysis framework for understanding the mechanistic principles that generate large-scale patterns of neural activity. In a large-scale biophysical model (LSBM) the nodes represent the electrophysiological dynamics of ensembles of neurons. These ensembles of neurons interact via neural white matter fibers (edges) that can be derived from diffusion tractography images [11–13]. Currently it is computationally prohibitive to simulate all neurons and synapses in the brain. Moreover, even if it were possible, modeling such a large number of elements would yield results that would be challenging to interpret. Therefore, the neural activity in an ensemble of adjacent neurons is reduced mathematically to a few macroscopic variables. These macroscopic variables are subsequently coupled through the network of white matter fibers and used to simulate neuroimaging data such as magnetoencephalography (MEG) or functional magnetic resonance imaging.

LSBMs have been used among others to determine the relationship between the anatomical structure of neural fibers, the connectome, and the functional connectivity (FC) observed during the resting state [11,14,15]; to assess the influence of source reconstruction or tractography seeding methods on neuroimaging analysis [16,17]; to create personalized models of seizure activity in epileptic patients or models aiding surgical interventions [18–20]; to analyze the sources and sinks of brain waves [21]; to model flows of information through the brain [22]; and to derive the conditions necessary for selective synchronization between ensembles of neurons [23].

However, the assumptions that are made to be able to derive the macroscopic variables which describe the neural activity in an ensemble of neurons impose limitations. A fundamental limitation is that the macroscopic variables cannot describe within-ensemble modulations of synchronization [5,12,24]. Yet, it is exactly these fluctuations of local synchrony (often referred to as event-related synchronization/desynchronization) that characterize behavioral tasks [25], and are associated with changes in functional connectivity between brain regions during tasks and on-going activity [3,4]. In addition, there is increasing evidence that these local fluctuations can be short-lived (i.e., transient and bursting) in tasks and resting-state [9,10]. Another limitation is more practical as the number of parameters often scales with the level of biological detail which makes it difficult to fit such models. As a result, most studies assume identical parameters for all ensembles, and thereby neglect between-ensemble differences.

While biologically realistic LSBMs are impractical to model between-ensemble differences, such differences have been modeled with non-biological models. The latter LSBMs model each ensemble as a Stuart-Landau oscillator [26,27]. Although one Stuart-Landau oscillator can describe the mean firing rate in an ensemble of neurons for particular parameterizations [28,29], this oscillator model is obtained from models of macroscopic activity that cannot capture modulations of local synchrony [12,24]. Moreover, when a set of Stuart-Landau oscillators with additive noise and various bifurcation parameters are coupled in a heterogeneous network with time-delays – as they are in LSBMs [27,30], it is very difficult to estimate the range of amplitudes of these oscillators. Therefore, it is hard to interpret their amplitudes as the local degree of synchrony and to compare amplitude differences across oscillators.

To obtain a LSBM that explicitly accounts for modulations of local synchrony and still has a low number of parameters, we introduce here a low-dimensional LSBM derived from a network-of-networks of Kuramoto oscillators. The Kuramoto oscillators are a canonical model of synchronization in biological systems, and they accounts for many of the dynamics of synchrony found in neural populations such as traveling waves and metastability [15,31,32]. Each sub-network of Kuramoto oscillators in this LSBM represents an ensemble of neurons within a particular cortical region, and its synchrony is given by the Kuramoto order parameter (KOP) [33,34]. The KOP has been found to be a good measure of synchrony in an ensemble of neurons whose dynamics are reduced with a mean-field approach [24,35,36]. In turn this mean-field reduction captures modulations of synchrony and explains event-related de/synchronization [5].

There are previous LSBMs that have used a network-of-networks of Kuramoto oscillators structure [20,23,37,38]. Yet, these LSBMs did not model resting-state FC, and their formulation is computationally expensive for LSBMs. Moreover their dynamics are influenced by the number of oscillators in the sub-networks and their natural frequencies [39,40]. Such finite-size effects are relevant for modeling resting-state because resting-state FC emerges at the point with the largest finite-size effects – the edge between asynchrony and partial synchrony [11,27,41]. To solve these problems we applied a mean-field reduction over a model with infinite oscillators on each sub-network. This reduction gave one equation per sub-network that describes the evolution of the local synchrony with the KOP. Moreover, the dynamics of the KOP can be manipulated with the local coupling parameter that represents the local connectivity strength.

Our LSBM simulated resting-state alpha band static FC (sFC) and time-resolved FC (trFC) of amplitude envelopes in two scenarios of increasing complexity. The first scenario assumes identical ensembles (homogeneous ensembles), whereas the second scenario generalized this to the case with different local connectivity strengths for each ensemble (heterogeneous ensembles). Our results show that FC emerges in both scenarios when high metastable dynamics (i.e., temporal fluctuations of synchrony) coexist within-ensembles and between-ensembles. At this working point, repulsion from local synchrony along with time-delayed attraction to global synchrony leads to coordinated fluctuations of local synchrony that are responsible for creating changing patterns of FC. At the same time, there are ensembles that are attracted to local synchrony which do not have FC, but influence other ensembles. An exception to this behavior are ensembles that represent parts of the default-mode network as they are attracted to local synchrony and also are functionally connected to each other.

## Results

### Dynamics of the large-scale brain model

The LSBM that we propose is defined by equations 1a and 1b, which describe the temporal evolution of the KOP in one ensemble (i.e. the local synchrony within the ensemble). The dynamics of this LSBM are governed by the global coupling parameter *G*, the local coupling of each ensemble *L*_*n*_, the spike-propagation velocity (proportional to the time-delays *τ*_*np*_), and the probability distribution of natural frequencies in each ensemble (*Ω*_*n*_ central natural frequency; *Δ*_*n*_ spread of natural frequencies). We focus on the impact of global and local couplings as well as time delays, and assume an identical frequency distributions for all ensembles (*Ω = 10.5 Hz, Δ=1*). The section *model and methods* explains this LSBM in detail.

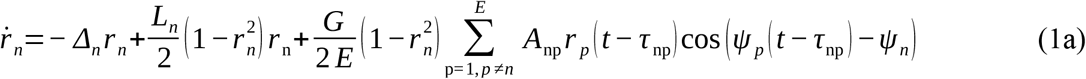

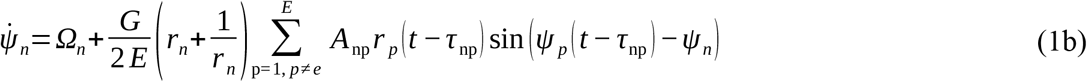

In these equations, the over dot represents the time derivative. Notice that we have removed the time dependency of *r* and *Ψ* when they do not have time delays. Equation 1a represents the temporal evolution of the level of synchronization within ensemble *n* (brain region). The variable *r* is bounded between zero and one, where zero means full desynchronization and one means full synchronization. Equation 1a can be divided into three parts by the plus signs. The first and second parts represent local dynamics, while the third part encapsulates the interaction with other ensembles.

- The first part opposes within-ensemble synchrony due to the heterogeneity of natural frequencies *Δ*_*n*_.
- The second part contains the local coupling parameter *L*_*n*_, which tunes the connectivity strength within the ensemble.
- The last part scales between-ensemble coupling strength, *G*, as well as the interaction delays, *τ*_*np*_. The interaction between ensembles depends on their phase differences and the local synchronies.

The contribution of *L*_*n*_ over *r*_*n*_ can be assessed by assuming that the global coupling is equal to zero. With this assumption, equation 1a becomes the mean-field reduction of a canonical Kuramoto model [42]. For this equation, there is a critical coupling value (*L*^*c*^) at *L*_*n*_ = 2Δ_*n*_. Thus, ensembles with *L*_*n*_ > 2Δ_*n*_ tend to synchronize, while ensembles with *L*_*n*_<2Δ_*n*_ tend to become desynchronized. The lower *L*_*n*_ (including negative values), the more asynchronous the ensemble, *r*_*n*_ → 0 [43]. In addition, the further away *L*_*n*_ is from *L*^*c*^, the stronger the influence of *L*_*n*_ on *rn.* Equation 1b describes the dynamics of the mean phase of the oscillations within the ensemble *n*. The mean phase evolves at the pace of its natural frequency, *Ω*_*n*_, plus the interaction with other ensembles scaled by *G/E* and approximately the inverse of its own local synchrony level. Therefore, locally desynchronized ensembles (small *r**n***) are strongly influenced by other ensembles, while locally synchronized ensembles (*r*_***n***_ ~ 1) are almost not influenced by others.

In what follows, we refer to *r*_*n*_ as the local synchrony. The phase synchronization among all ensembles, *R,* is referred as global synchrony. Global synchrony was measured as the KOP of the local phases as follows,

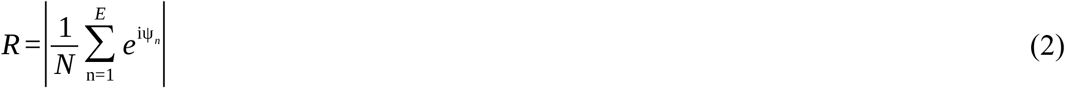

The standard deviation of the modulus of the KOP over time (i.e. *SD(abs(*KOP))_*t*_) is a measure of the metastability [44]. Therefore we describe the dynamics of the LSBM in terms of global metastability, *SD(R)*_*t*_, and local metastabilities, *SD(r*_*n*_)_*t*_.

### Scenario 1: Ensembles with homogeneous local couplings

In the first simulation scenario, we searched for optimal global parameters (global coupling, and spike-propagation velocity), while we held the local coupling identical for all ensembles (the local coupling will be optimized in the second scenario, during which the global parameters are kept at the levels determined by this first scenario). We assumed a constant spike-propagation velocity and time delays proportional to the Euclidean distance between nodes. The optimal parameters were found by two independent stochastic optimizers (particle swarm optimization, PSO; and adaptive differential evolution, aDE). The fitness function of the optimizers maximized the correlation between simulated and MEG sFC in the alpha band, while it was constrained to biologically plausible solutions (see sections *Fitness function*, and *Optimization constraints*). sFC was measured as the correlations of the low-pass filtered ampliude envelops of the analytic alpha band signals. The optimal solutions had a correlation of ~0.55 between simulated and MEG sFC (Figure 1). The optimal global parameters of the two optimizers were identical up to the second decimal point.

**Figure 1.**
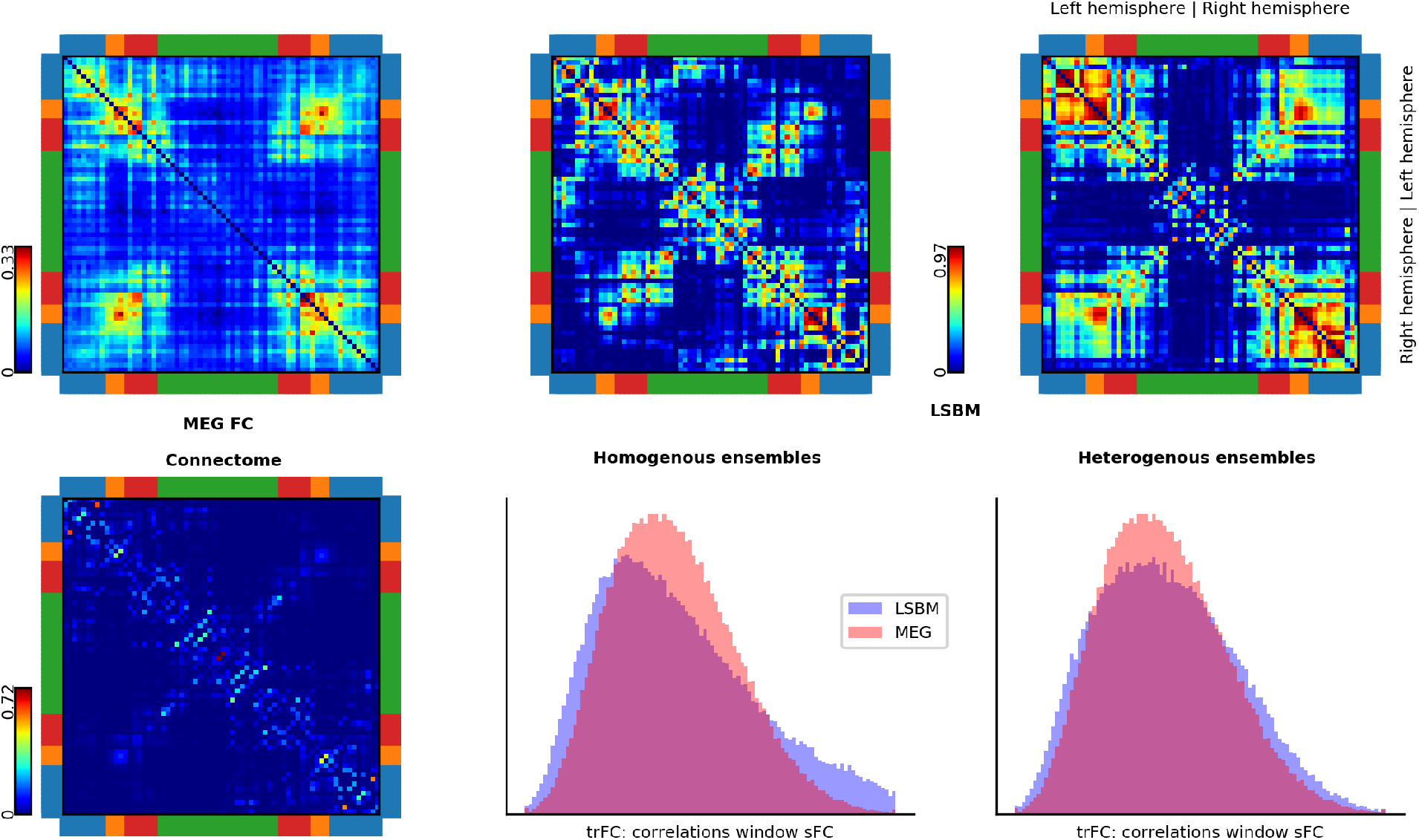
Connectome and FC from real MEG data and from simulations. First column shows resting-state MEG static FC (orthogonalized alpha band envelope correlations, top), and the anatomical network or connectome (bottom). The second and third columns show the FCs obtained from simulations with homogeneous and heterogeneous ensembles, respectively. The top row shows the static FCs. The bottom row shows the histograms of time-resolved FC recurrence (trFC: correlation of static FCs for 15 sec. moving window with 12 sec. overlap) for simulation (blue) and MEG data (pink). Brain lobes are color-coded around the connectivity matrices – blue, temporal; orange, occipital; red, parietal; and green, frontal.

Figure 2 shows that FC emerges when the local coupling is below the critical coupling, *L*^*c*^. Being below *L*^*c*^ would lead to asynchronous ensembles if they were decoupled from the rest of the brain. However, when there are between-ensemble interactions, the local synchrony can increase. Suitable local and global couplings have a negative correlation. The global coupling increases as the local coupling decreases. Moreover, Figure 2 shows that time-delays are needed to reproduce sFC.

**Figure 2.**
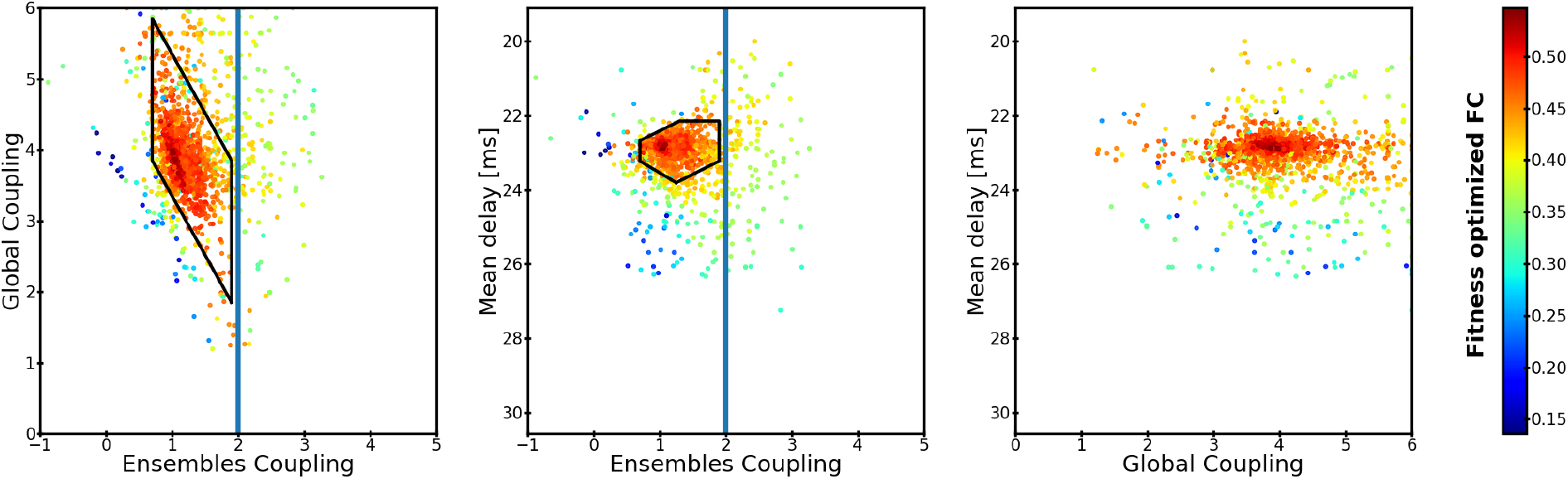
Similarity between simulated and MEG sFC during the optimization with homogeneous ensembles. Each dot represents the correlation between simulated and empirical MEG sFC during one evaluation of the fitness function. A three-dimensional parameter space consisting of global coupling, local coupling, and mean delay (proportional to spike-conductance velocity) is optimized for reproduction of the empirical sFC. Each panel has one dimension collapsed over the two axes. Simulations outside of the optimization constraints are not shown. The black area indicates the area of the parameter space that is further analyzed for trFC. Only the parameter combinations which produced dynamics within the biological constrains are shown.

A LSBM parametrized within the same range but without time delays leads to full global synchrony (R ~ 1) and steady partial local synchrony (0 < *r*_***n***_ = constant < 1) proportional to the coupling strength with other ensembles. On the contrary, a LSBM with long time delays becomes globally asynchronous with steady partial local synchrony. This is consistent with previous results obtained analytically in a model with homogeneous couplings [45].

Global and local metastability are relatively high, but not maximal, in the area that better predicts FC (see Sup. Fig. 2). The highest global metastability appears to be above *L*^*c*^, although the simulated sFC has low similarity with MEG sFC. The highest local metastability appears below *L*^*c*^. In other regions of the parameter space both metastabilities tend to be lower than in the area that reproduces FC better. The averaged global and local synchronies are moderately high and almost constant in the area that best predicts FC. Outside of this area, global and local synchrony either increases or drops abruptly (see Sup. Fig. 2).

Next, we evaluated the trFC within the area of the parameter space that reproduces sFC (black polyhedrons in Fig. 2). trFC was measured as the recurrence (Pearson correlation) of sFCs over a 15-second sliding window with 13 seconds overlap. Histograms of recurrence values were built for both simulated and MEG data. The similarity between these two histograms was measured with the Kolmogorov-Smirnov distance (KS-distance). The parameters that best predicted 300-second sFC and trFC simultaneously were associated with the highest concurrent local and global metastabilities (Figure 3). The best trFC had a KS-distance of ~0.13, and the best sFC had a correlation of ~0.52. The Supplementary Figure 3 shows the trFC KS-distances and sFC correlations over the polyhedron in Figure 3. The global parameters that provided the best joint fit of sFC and trFC were used in the second scenario (this corresponded to: spike-conductance velocity ≈ 3.42 m/s, and global coupling ≈ 3.57; magenta arrow in Figure 4).

**Figure 3.**
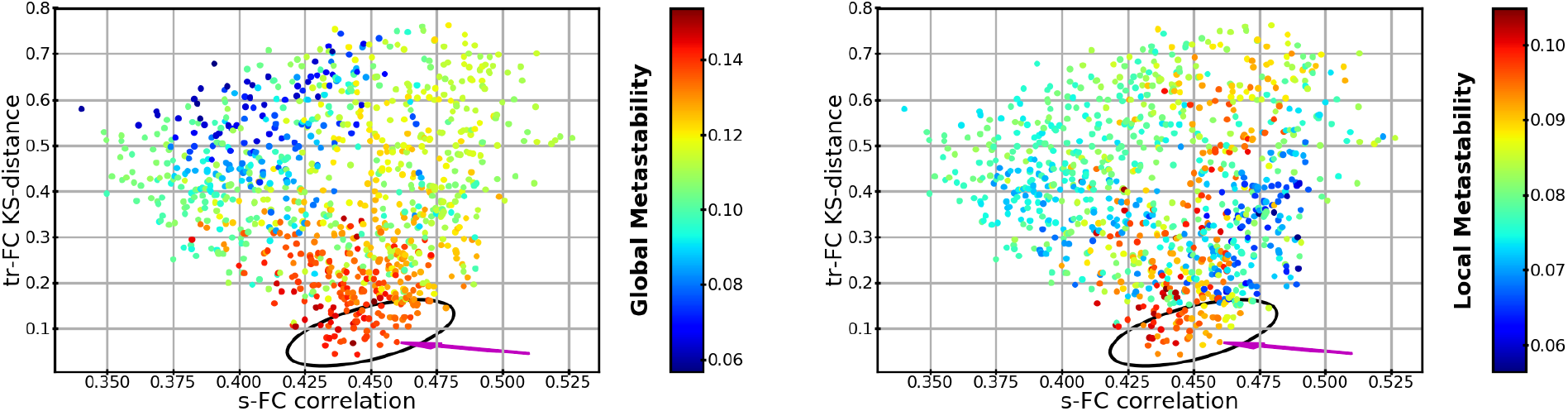
Global and local metastability as a function of trFC and sFC similarity between 300-second simulations and MEG. Each dot has the same conventions as Figure 4, but the dots are arranged by similarity to trFC (KS-distance; y-axis), and similarity to sFC (correlation; x-axis). (Left panel) Dots are colored by their global metastability. (Right panel) Dots are colored by the local metastability averaged over ensembles. The black ellipsoid marks the areas that provide a good compromise between trFC and sFC fit. Magenta arrows indicate the simulation from which we took the global parameters for the second scenario.

**Figure 4.**
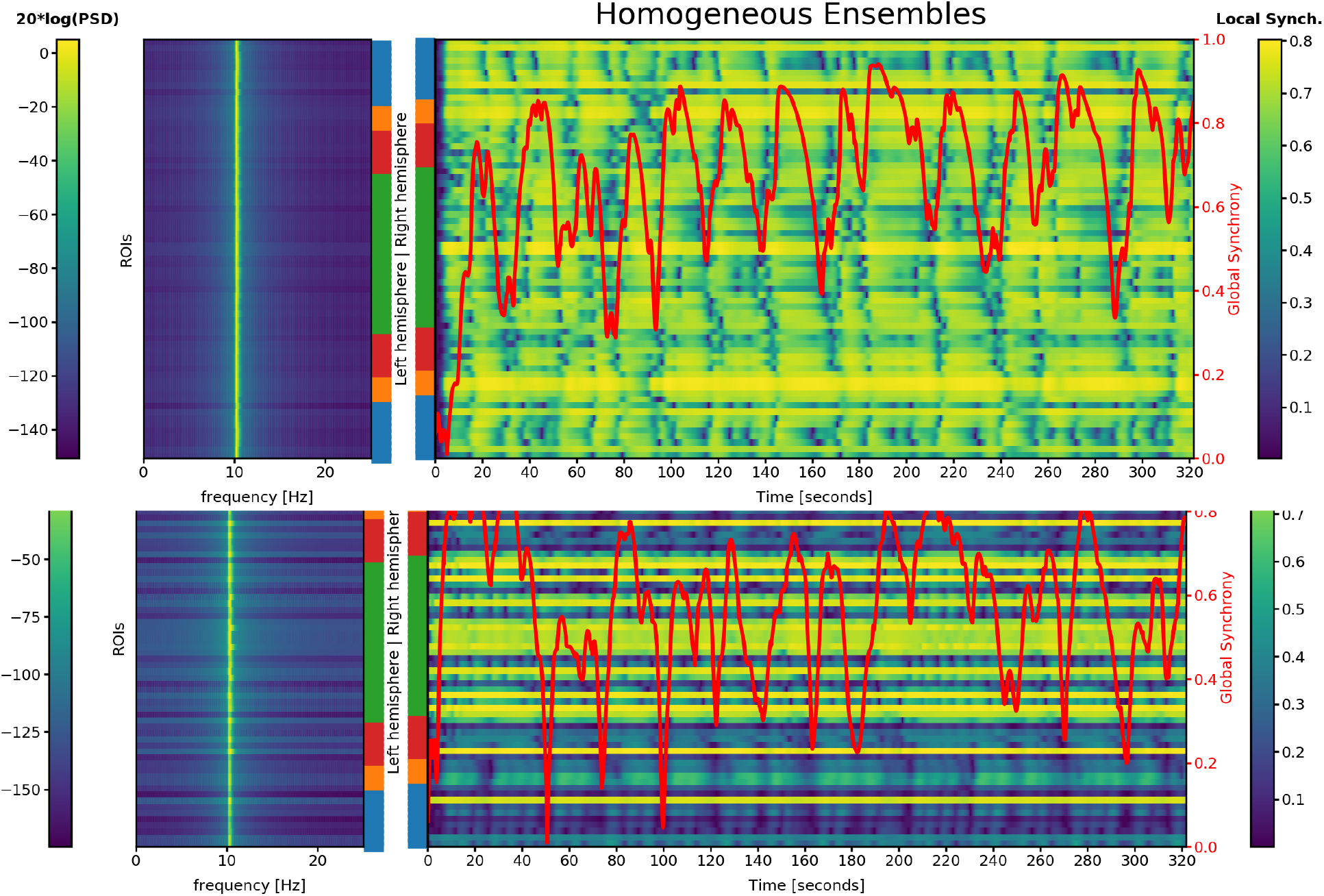
Power spectral density (PSD) and temporal evolution of local synchrony in each ensemble, overlaid with the level of global synchrony. Top: LSBM with homogeneous ensembles; Bottom: LSBM with heterogeneous ensembles. Each horizontal line reflects one brain region. The lobes are color-coded using the same conventions as previous figures. The left panel shows the power spectrum of the central frequency from the probability density of frequencies. The right panel shows the temporal evolution of local (within-ensemble) synchrony on yellow-blue color. The global synchrony (between-ensembles) is represented in red (right, y-axis).

A simulation with a good fit to MEG (Figure 3; magenta arrow) is shown in Figures 1 (sFC and trFC) and 4 (time-courses of synchrony and power spectra). The power spectra shows the central frequency of each ensemble. However, the power density at the microscopic level would be broader than in Figure 4 (except if *r*_*n*_ = 1) because the LSBM reduces a probability density of frequencies to its mean-field frequency. It is a non-trivial task to obtain the individual frequencies given the mean-field frequency. Local synchrony fluctuates at different rates at each ensemble. These fluctuations are not strictly periodic because they do not have a constant frequency. Nevertheless some regularities are visible at different time scales within and between ensembles. The global synchrony has large fluctuations as well. These aperiodic but temporally structured fluctuations at both spatial scales resemble metastable dynamics. The first 20 seconds of simulation correspond to the initial transient dynamics that were not included in the FC.

### Scenario 2: Ensembles with heterogeneous local couplings

Having established the global parameters of our LSBM, we then turn to the scenario in which local couplings can differ between ensembles. Here, the optimizers identified the local couplings that reproduced sFC, while the global parameters were kept constant (derived from the first scenario). To reduce the dimensionality of the parameter space, we assumed equal local couplings in homotopic ensembles as the sFC and the anatomical networks are almost symmetric respect to the interhemispheric fissure.

PSO and aDE achieved a maximal correlation between simulated and MEG sFC of 0.80 and 0.78, respectively. The optimal parameters found by each optimizer were not identical (0.81 correlation & 0.84 cosine similarity), and neither were the sFCs generated (0.90 correlation & 0.94 cosine similarity). Supplementary Figure 3 shows the local coupling parameters used in each optimization iteration sorted by the correlation with MEG sFC. Next, we looked at the 300-second sFC and trFC produced by the 1000 best solutions from each optimizer.

Figure 5 shows that the solutions with high similarity to MEG data tend to have high global and local metastabilities, although there are some differences in metastability for solutions with comparable fit to sFC and trFC (see later). The best fit to MEG trFC has a KS-distance of ~0.02, and the best correlation to MEG sFC is ~0.79.

**Figure 5.**
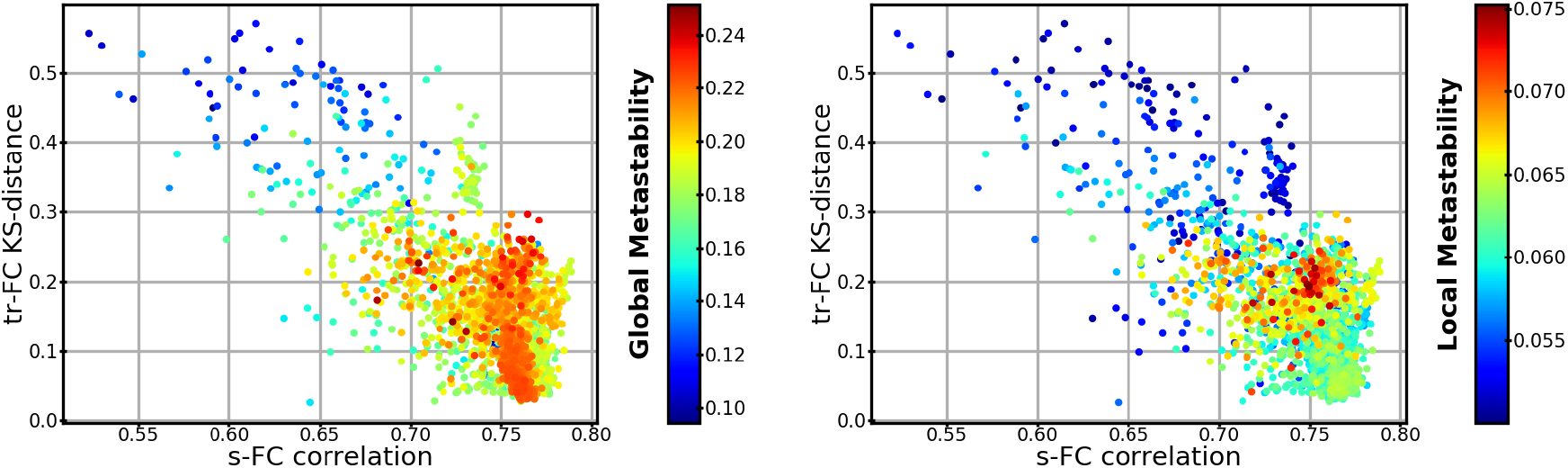
Global and local metastability of heterogeneous ensembles as a function of trFC and sFC similarity between 300-second simulations and MEG for the best 2000 simulations. Each dot represents one simulation similar to Figure 3. (Left panel) Dots are colored by their global metastability. (Right panel) Dots are colored by their local metastability.

Figure 6 shows in red the local couplings for the solutions that achieved the best compromise between sFC and trFC similarity to MEG data (correlation > 0.75 and KS-distance < 0.04), a set of 49 solutions. There is not a unique combination of local couplings which predicts sFC and trFC. Some ensembles—such as the superior temporal or the inferior temporal regions—can take a wide range of local couplings, while others—such as the precuneus—work within a narrow range of local couplings. There are other ensembles like the parahippocampal area, the cuneus, or the posterior cingulate that can take local couplings from two disjoint sets of values. For example, the local couplings at the posterior cingulate group either around 6 or −3. Next, the local couplings at the posterior cingulate region were used to divide these 49 solutions into two groups. The group with local couplings above 2 is shown with the orange histogram in Figure 6. The solutions from each group had the local couplings reconfigured in a way that produced almost the same sFC and trFC.

**Figure 6.**
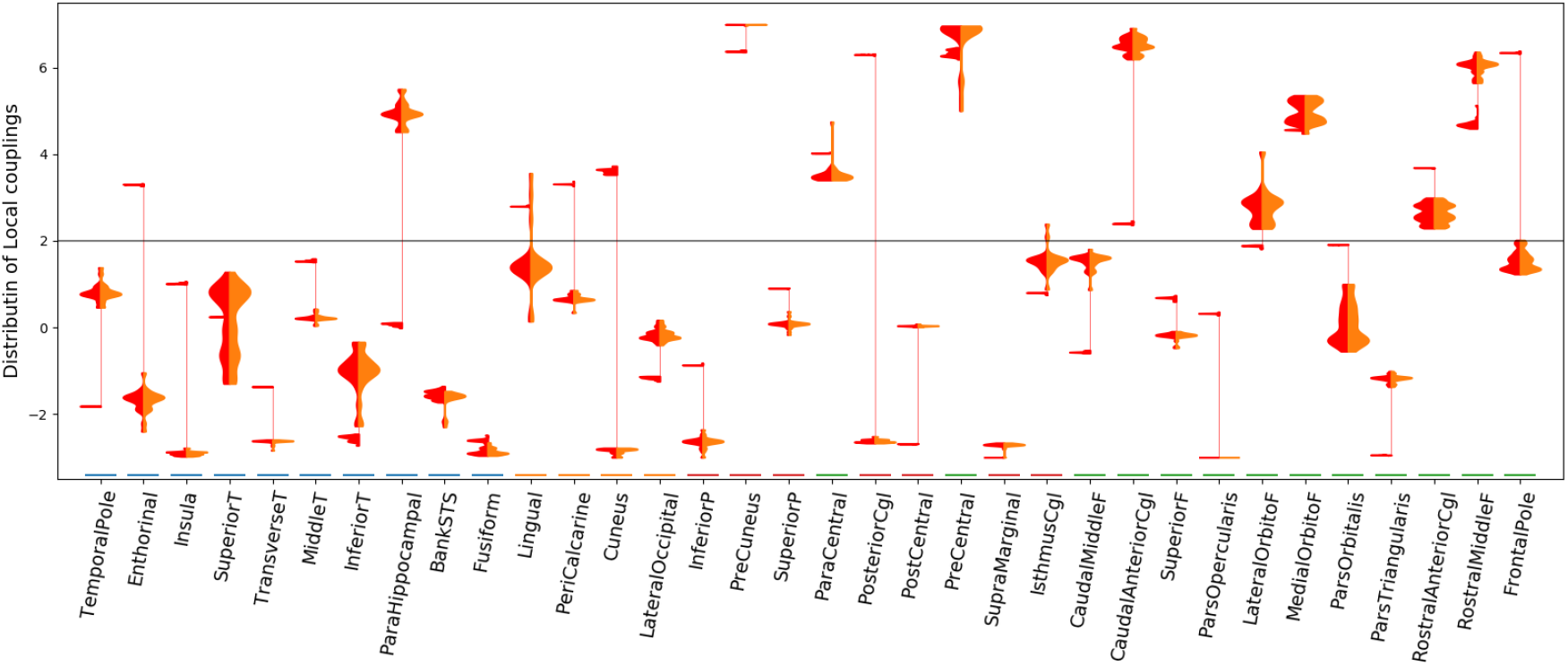
Density estimates of local couplings by brain regions for the best solutions in terms of sFC and trFC similarity. Each column on the x-axis is a brain region. In each column, there are vertically two density estimates of local couplings with the value indicated by the y-axis. The densities on the left (red) contain the number of solutions that best predict sFC and trFC for a given local coupling (y-axis). The vertical thin red line connects clusters of solutions in the same brain region. The densities on the right (orange) are a subset of the local couplings of the densities on the left. This subset contains solutions from only one cluster of local couplings. The horizontal gray line indicates the critical local coupling.

The global metastability is significantly different (Mann-Whitney U test) between the groups separated by local the coupling at the posterior cingulate (p-value < 10^−8^) as well as the local metastability averaged over ensembles (p-value < 10^−6^). Such differences in metastability are also significant for solutions with sFC correlation higher than 0.7 and trFC KS-distance lower than 0.15. These differences are noticeable in Figure 6 as well.

Next, simulated local metastability was compared with local MEG metastability. Local MEG metastability for a particular brain region measured as the standard deviation over time of the envelope from the Hilbert transformed alpha band signals. Local metastability was measured in each participant independently, and subsequently averaged across participants. The group of solutions depicted in orange in Figure 6 had a median correlation of 0.13 between MEG and simulated local metastability, while the other group of solutions had a median correlation of −0.14. This shows that although FC is very similar for both groups of local couplings, the metastable dynamics can be quite different. In what follows, we will show only results from the cluster of local couplings which have a positive correlations with MEG data (orange histogram in Fig. 6).

The local couplings have a significant negative correlation with the simulated nodal sFC strength (sFCS; average correlation of −0.46 and −0.40 for the right and left hemispheres respectively, p-values < 10^−5^). The nodal sFCS is the sum of sFC in a node, so brain regions with high functional connectivity to other regions have high sFCS. Figure 7 shows that there is a general tendency of areas with high sFCS to have a local couplings below the critical local coupling, *L*^*c*^. In contrast, areas with lower sFCS have a heterogeneous arrangement of local couplings dominated by couplings above *L*^*c*^. There are a few exceptions to this, such as the precuneus or the parahipocampal which have relatively high sFCS and high local couplings, or the insula and the parsorbitalis which both have low local coupling and low sFCS. Simulated and MEG sFCS follows the same pattern except for some asymmetries between hemispheres. For example, the simulations show a strong sFCS asymmetry in the precuneus, and the isthmus cingulate which are not present in the MEG data. The local couplings were not correlated with the nodal connectivity strength of the anatomical network (*ρ* < 0.07 in right and left hemispheres).

**Figure 7.**
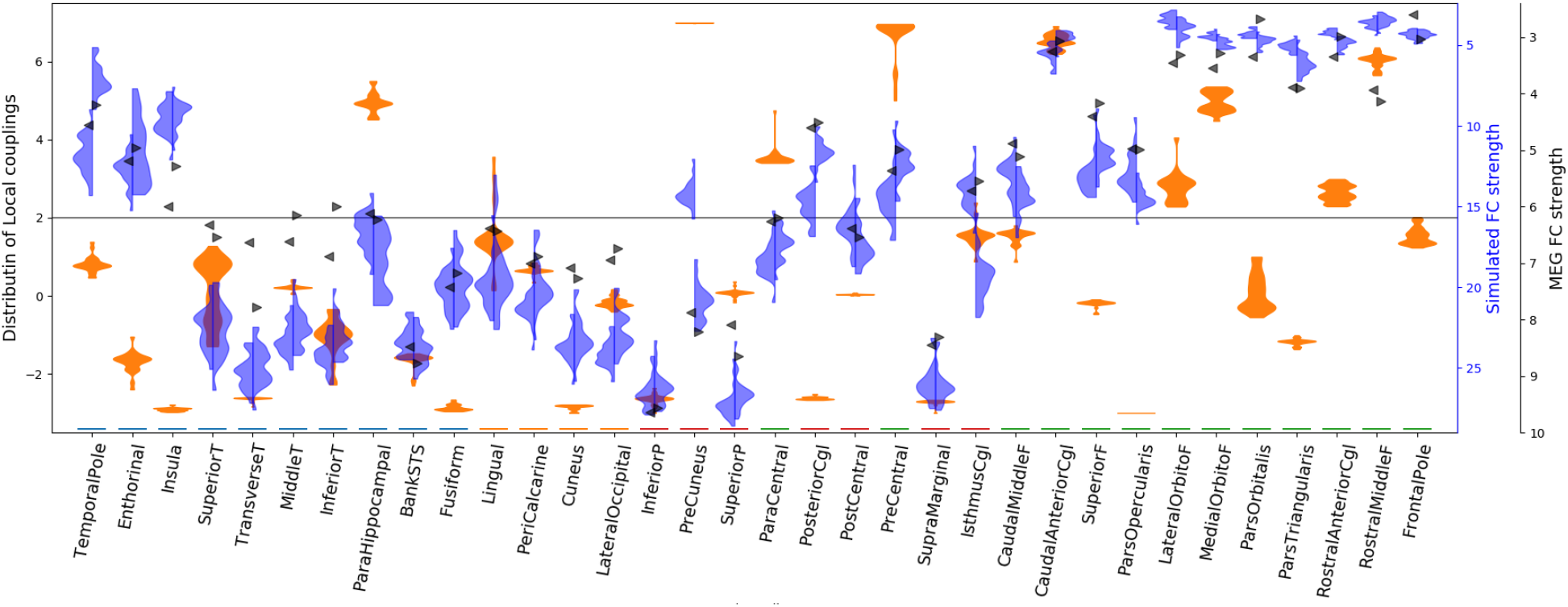
Density estimates of local couplings and functional connectivity strength by brain region. Each column on the x-axis is a brain region. In each column there are: one two-sided density in orange, two one-sided densities in blue (left/right), and two arrows in black. The orange density contains the same local couplings that are shown in orange in Figure 6. The magnitude of local coupling is indicated on the left y-axis. Blue densities represent the sFCS for these local couplings, left and right for each hemisphere. The black triangles indicate the MEG mean sFCS across subjects for each hemisphere. Both scales of FCS are on the right y-axis with the respective colors. The scales of sFCS are inverted to aid visual comparison with local couplings as they have a negative correlation (−0.46 and −0.40 for the right and left hemispheres respectively). The horizontal gray line indicates the critical local coupling.

The sFC and trFC of one these solution are shown in Figure 1, and the associated power spectra as well as the synchrony time courses are shown in Figure 4. Similar to the heterogeneous ensembles, there is a variety of short- and long lasting patterns of within- and between-ensembles synchrony that resemble metastable dynamics. However, in the model with heterogeneous local couplings there is higher variability of local and global synchrony than in the mode with homogeneous local couplings. The Supplementary Figure 7 shows the spectrogram of the solution shown in Figures 1 and 4.

## Discussion

We have derived a parsimonious large-scale brain model (LSBM) that is able to successfully simulate the dynamics of resting-state MEG functional connectivity. This LSBM reconciles simplicity with biological interpretability and allows for the manipulation of both local and global neural synchrony within one framework. Such modulations of synchrony are observed in neuroimaging data at multiple spatial and temporal scales, and are believed to be fundamental property of neural activity [1–3,5–8,18,25,27,46] Our model goes beyond traditional LSBMs that are not able to capture modulations of local synchrony [5,12,24]. Moreover, the tendency of each ensemble to de/synchronize can be adjusted with a single parameter that represents within-ensemble connectivity strength. Because local synchrony can be manipulated with one parameter, it is feasible to fit LSBMs with heterogeneous local synchronies and to analyse between-ensemble differences.

The proposed LSBM was able to simulate resting-state sFC and trFC of MEG alpha band envelopes in two scenarios. In the first scenario all ensembles were identical (homogeneous ensembles). In the other scenario each ensemble could take a different local coupling parameters (heterogeneous ensembles).

### Scenario 1: Ensembles with homogeneous local couplings

The LSBM with identical ensembles showed that FC emerged when there were interaction delays between ensembles. This is in agreement with findings of previous LSBMs that relied on different reductions of neural activity [14,16,30,47]. Such interaction delays between ensembles of neurons are due to finite spike-conductance velocities along neural fibers [48,49].

Another consistent finding in LSBMs is that FC emerges when the dynamics of the ensembles are close to a critical point. Below this critical point, ensembles do not oscillate, but once the critical point is exceeded, the ensembles start to oscillate [11,26,27,41]. For example, a stability analysis of LSBMs with several reductions of neural activity showed that additive noise as well as delayed coupling with other ensembles induced oscillations and plausible patterns of resting-state sFC [41]; LSBMs based on Stuart-Landau oscillators reproduced sFC and trFC only when the ensembles were near a supercritical Hopf bifurcation [26,27]. Similarly, FC emerged in our LSBM if the local coupling was just below a critical local coupling, *L*^*c*^. *L*^*c*^ was the minimum coupling required by an unperturbed ensemble to leave the desynchronized state. Below *L*^*c*^ the ensembles pushed towards local asynchrony while local couplings above *L*^*c*^ promoted local synchronization.

Ensembles that had low local coupling, reflecting desynchronization, were more sensitive to the phase of other ensembles, which facilitated between-ensemble synchrony. This increase of between-ensembles synchrony raised local synchrony, but local synchrony could not be maintained as between-ensemble synchrony was perturbed by interaction delays, and there was not enough local connectivity to sustain the synchrony from within the ensemble itself. Therefore, the ensembles went into a state of fluctuating partial local synchronization that gave rise to FC networks.

We have shown that FC depends on a balanced interaction between the local coupling and the global coupling among ensembles. When the global coupling increased, the local coupling decreased proportionally. However, if only the global coupling increased, the LSBM became fully synchronized globally and locally – a pathological state that occurs during epileptic seizures [50]. When only the local coupling decreased, the ensembles became asynchronous and the fluctuations of local and global synchrony stopped. In other words, in order to maintain the state of fluctuating partial synchrony responsible for FC, the ensembles counterbalanced an increase of attraction to global synchrony by decreasing local synchrony and vice versa. Hence, FC emerged from precise opposition between local attraction to asynchrony and between-ensembles attraction to synchronicity, the latter being perturbed by interaction delays. This relationship between local and global coupling suggest that the critical local coupling was shifted towards lower local couplings if the global coupling increased. Other LSBM have included a local feedback inhibitory control mechanism to counteract the increase of global coupling [51,52]. Feedback inhibitory control adapts the strength of recurrent inhibitory synapses to compensate for the increase in excitatory activity due to long-range connections with other ensembles. Feedback inhibitory control increased the predictability of FC for spontaneous and evoked activity [51,52].

Interestingly, FC was reproduced when high metastability (fluctuations of synchrony) coexisted at global and local scales. Metastability is a hallmark of brain dynamics and it occurs at multiple spatial scales. Metastability allows to simultaneously integrate and segregate information [21,46]. Metastable dynamics are believed to provide the neural flexibility needed to adapt and respond fast as well as to maintain current states like memories [46,53]. Because previous LSBM could not capture fluctuations of local synchrony [5,12,24], they reported only high global metastability without corresponding local metastability [15,26,27]. In contrast, our LSBM showed maximal global metastability when local metastability was low and sFC was poorly reproduced. This suggest that accounting for fluctuations of local synchrony might be necessary for successfully modeling large-scale brain networks.

### Scenario 2: Ensembles with heterogeneous local couplings

The LSBM with heterogeneous ensembles allowed for ensembles which could promote local synchrony or aynchrony at various levels. This model simulated FC with higher accuracy than the model with homogeneous ensembles. This was not entirely surprising given that the model with heterogeneous ensembles had 34 parameters to tune local dynamics compared to a single parameter in the model with homogeneous ensembles.

More interestingly, the plausible local couplings of each ensemble exhibited different distributions, including bimodal distributions. This suggest that there is not a single way to generate a particular target FC, but instead it can be generated by multiple different configurations of local dynamics. Such pattern points at an adaptive property of the brain, which can function in a number of different states, thereby making it resilient to perturbations. A similar effect is observed at a lower scale as the same macroscopic activity can be obtained form a network of neurons with different synaptic strengths [54].

The local coupling had a negative correlation with the sum of nodal sFC strength (sFCS). In other words, ensembles with lower local couplings (below *L*^*c*^) had a tendency to higher sFCS, and vice versa. Similar to the LSBMs with homogeneous ensembles, the balance between the local tendency towards asynchrony and the time-delayed between-ensemble synchrony gave rise to fluctuations of local synchrony and FC. In contrast, ensembles with high local couplings (above *L*^*c*^) had a tendency to synchronize locally and did not engage into coordinated fluctuations of local synchrony, i.e. FC. Such ensembles were more attracted to local than between-ensemble synchrony. Nevertheless, they had strong influence on other ensembles driving ensembles with low local coupling. Some exceptions to this observation were ensembles that belong to the default-mode network, such as the precuneus or the parahipocampal areas [55], which tended to have high local couplings and relatively high FC as well. Such high local coupling may reflect a state of high local coupling and low excitability that allows for maintaining the local dynamics in the face of external disturbance. Future research should investigate these individual differences in dynamics between the different ensembles.

In the simulations there were ensembles with asymmetric sFCS between hemispheres which were different from the MEG sFCS asymmetries (e.g. the precuneus). This discrepancies might be due to the assumption of equal local coupling at homotopic brain regions. To our knowledge there is only one study so far which has attempted to optimize all cortical brain regions in a LSBM [27]. This study showed that some homotopic brain regions had different parameters. However it is not possible to compare parameters systematically across studies because the topologies of the FC are considerably different and the neuroimaging technologies are different (FC in [27] included frontal but not occipital regions and it was measured with fMRI). Although the assumption of homotopy reduced our ability to fit the functional data, we decided that this was the best trade-off between model complexity and model fit.

### Optimization framework

While we have argued that a multimodal distribution of plausible local couplings can have reasonable biological underpinnings, there exist other plausible explanations. First, it could be that the optimizers explored two local optima, but they failed to find a global optimum. It is known that stochastic optimization do not guarantee finding a global optimum. However, we think this explanation is unlikely because we have obtained similar results in multiple runs of different optimizers. It is therefore likely that there were two global optima.

Second, we used sFC and trFC to assess the similarity between simulations and MEG data. The fitness function of the optimizers only evaluated the similarity of sFC, but this was sufficient to predict trFC with high precision. However local and global metastability were different among the two clusters of local couplings. Adding local information to the fitness function might help to constrain the solutions of the optimization. Here we focused on optimizing FC as this is the most common approach on the literature. Moreover, we find relevant the fact that similar FC could emerge from different local dynamics. This might contribute to explain the variability in neuroimaging measurements between and within-subjects during-resting sate as well as behavioral tasks.

Third, not all optimizers were able to find suitable solutions. For example a successful optimizer on similar problems (covariance matrix adaptation evolutionary strategy [56]) was not able to find proper solutions in this context, while aDE and PSO worked satisfactorily. aDE and PSO found similar solutions, although they relied on different heuristics. These heuristics were noticeable in their evolution towards the best solution. PSO was more exploitative, while aDE was more explorative. PSO found a good set of parameters fast, and then restricted exploration near these parameters. On the contrary, aDE favored diverse combinations of parameters, but took longer to find a good set of parameters. Given the dissimilar behavior of the two optimizers, a further improvement could be to try other algorithms or to migrate solutions between optimizers. Strategic migrations of parameters tends to improve the optimization[57]. However, this enhancement might not avoid converging to local optima, nor may it improve the similarity between simulated and MEG data. Recently an optimization approach based on Bayesian Gaussian-process optimization gave remarkable results in a LSBM with a 5-dimensional-parameter space with less computing time [16]. Further research on this framework with higher-dimensional-parameter spaces could fuel the usability of LSBM.

### Limitations and extensions

In this LSBM we made several assumptions and choices that might influence the results. First, we imposed a network-of-networks structure where the higher level nodes represent ensembles of neurons in each region of a cortical parcellation. The parcellation scheme determines the number of brain regions, their size, and the topology of the anatomical network that connects the ensembles. This should be taken into account when interpreting or comparing the results with other studies. Moreover sub-cortical structures were not included, although synaptic noise in the thalamus has been related with multistable changes of alpha band amplitude [58]. The study of such noise-driven changes of amplitude was limited to a single region, but sub-cortical structures might be relevant for faster changes on FC and enhanced metastability.

To derive the model we assumed that each ensemble is a fully connected network. This assumption could be relaxed. A similar model to ours that contained random Erdös-Réniy topologies at the ensembles showed analytically and numerically that the same synchrony states are achieved with a shift in the coupling parameters proportional to the average degree of the networks [59]. Manipulating the topology of the connectivity of the ensembles might be useful to understand its impact on FC [60]. Moreover topological manipulations could contribute to understanding the effects of neurological disorders and different kinds of brain damage arising from natural causes or surgical interventions [19,60]. For example, LSBMs with a network-of-networks of Kuramoto oscillators have previously been used to understand abnormal neural synchrony [20,37,38]. These studies found, for example, that hyper-synchronous activity was found to emerge more easily from structurally central regions and when neural fibers had altered properties [38]. In addition, they demonstrated that hyper-synchrony can emerge more easily form the resting-state FC of epileptic patients [20,37], although these studies used the network of FC to couple the ensembles instead of the network of neural fibers. A network-of-networks of Kuramoto oscillators is an effective method to derive analytically the features of the brain that allow specific patterns of FC [23].

Another local heterogeneity that could be introduced is a diversity of density distributions of natural frequencies. We assumed that all ensembles have identical distributions of natural frequencies. Nevertheless heterogeneous natural frequencies might be a relevant factor because it has been observed in EEG data that the alpha peak frequency differs between brain regions during resting-state and behavioral tasks [61]. Such difference infrequencies might perturb global phase synchronization on the same manner as time-delays.

One may argue that a limitation or virtue of this LSBM is that it models only one frequency band but not the whole power spectrum of MEG. Yet, MEG is typically analyzed in frequency bands which each seem to play distinct functional roles [3,5,7,25,62–64]. During behavioral tasks as well as resting-state each frequency band tends to produce a different FC pattern [3,7,64]. Moreover, behavioral studies analyze event related synchronization within one frequency band [5,25,65]. Therefore, this LSBM provides a simple framework for modeling single-frequency-band electrophysiological data. Additionally the mathematical framework of the LSBM allows for multimodal distributions of frequencies [66]. Here we focused on reproducing FC in the alpha band, but we hypothesize that FC could be reproduced in other frequency bands simply by shifting the distribution of natural frequencies to the same frequency band, as a Hopf bifurcation model has already demonstrated successfully [26].

We have shown that the LSBM reproduces not only static FC, but also time-resolved FC. However, time-resolved FC might be influenced by the length of the moving windows used to estimate short-lived FC states. We chose a conservative length of 15 seconds, which may have missed short-lived FC patterns [7].

Finally, we assumed that all neural fibers have the same spike-propagation velocity, and that the length of the neural fibers is the Euclidean distance between ensembles. However there is a broad range of possible velocities which depend on the physiological properties of the neural fibers [48,49]. Some of this variability could be incorporated in the LSBM by using a probability density function of time delays [67]. Furthermore, other simulation studies have shown that the density of inter-hemispheric neural fibers is underestimated, and better predictions of FC can be achieved by scaling the inter-hemispheric connections [16,17]. Additionally, we assumed that homotopic ensembles have the same local coupling to exploit the symmetries on the anatomical and FC networks as well as to speed up computations. Even though our results suggest that without this assumption the similarity between MEG and simulations could improve, we think that this assumption is a good compromise between model complexity, computational time, and explanatory power.

### Significance

This study shows that resting-state MEG sFC and trFC emerge from the opposition between local asynchrony and global synchrony perturbed by interaction delays. Local asynchrony is facilitated by low local connectivity. Ensembles with low local connectivity adapt rapidly their phases to synchronize with other ensembles which suggest higher excitability. The increase of between-ensemble phase synchrony leads to higher local synchrony. However, between-ensemble synchrony is broken intermittently by the interaction delays, causing local synchrony to drop. These dynamics create coordinated fluctuations of local synchrony. The dynamics of the model are highly metastable at global and local scales when FC emerges. We also observed ensembles with high local connectivity which did not engage in FC, but still influenced the dynamics of ensembles that are responsible for FC. Parts of the default-mode networks show a differential behavior with high local coupling and strong FC.

We have presented a parsimonious large-scale brain model that balances simplicity and biological interpretability. Such a framework might contribute to determining the conditions and mechanisms that lead to the patterns observed in neuroimaging data. We have shown that this model can realistically simulate resting-state FC, and we hypothesize that it could be used to simulate changes in FC over a course of a behavioral task as well [3]. Moreover, our results suggest that modulations of local synchrony are fundamental for large-scale synchronization and FC. In the future we aim to use this model to understand which and how brain regions contribute to switch between transient functional connectivity networks during a behavioral task, which will contribute to elucidate how cognitive functions can arise from specific patterns of functional connectivity.

## Methods and Model

### Modeling one ensemble of neurons

The LSBM presented here aims at describing the evolution of the neural synchrony in an ensemble of neurons (cortical region) with a temporal resolution similar to MEG measurements. Moreover the LSBM should have the same framework for local and global synchrony. The model should have only one parameter to tune local synchrony, but it should allow for incorporating other properties of neural ensembles. The model is motivated by recent findings showing that modulations of synchrony in an ensemble of neurons can be expressed in terms of the Kuramoto order parameter (KOP) [5,24,35,36].

In our LSBM the neural activity within an ensemble of neurons is describe by the KOP, and the KOP is obtained form a network of Kuramoto oscillators [33,34]. Kuramoto oscillators can describe many synchronization phenomena observed on neural ensembles [31,32]. The KOP is a complex number with the modulus bounded between zero and one. When the modulus of the KOP, *r*_*n*_, is zero the neurons fire asynchronous, so the total electric potential produced is minimal. When the modulus of the KOP is equal to one, the neurons fire in a fully synchronized manner, resulting in a maximal electric potential.

### Modeling a network of ensembles of neurons

In a LSBM an ensembles of contiguous highly connected neurons is typically represented as a node in a fixed network whose edges are neural fibers. The set of edges connecting ensembles of neurons form a heterogeneous weighted multigraph. One weight of the multigraph is proportional to the density of neural fibers, *A.* The second weight of the multigraph represents the distance between the ensembles, *D*. This distance might impose time delays in the interaction between ensembles due to finite spike-conductance velocities [48,49]. At the ensemble level it is reasonable to assume that the interaction delays are negligible and the neurons are fully connected. In this way, the brain model becomes a network-of-networks. The section *Anatomical network of neural fibers* has more details about computing the multigraph of neural fibers.

The KOP for each ensemble of neurons was derived with a mean-field reduction in a network-of-networks of Kuramoto oscillators. To enforce such structure, the couplings within-ensemble were instantaneous, and identical, while the couplings between ensembles were delayed and weighted by the multigraph of neural fibers. In addition, the couplings within-ensemble were much larger than the couplings between ensembles. Kuramoto models with a network-of-networks structure have been studied previously [20,23,45,59]. We used a Kuramoto model and a mean-field reduction similar to the one analyzed by Skardal et al. [45,59]. The mean-field reduction is based on the Ott-Antonsen ansatz [42]. With this reduction the dynamics of each ensemble are given by the KOP (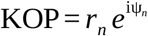). The temporal evolution of the KOP is dictated by equations 1a and 1b in the results section (S15a and S15b in the Supplementary information). In total the LSBM consists of *E* complex delayed differential equations where *E* represents the number of ensembles (brain regions). The Supplementary information provides all the mathematical details needed to derive the LSBM.

#### Model parameters

Some parameters of the LSBM can be given beforehand, while others should be estimated. The natural frequencies on the ensembles were set beforehand by a Lorentzian distribution function with center frequency *Ω* and spread *Δ.* We limited our simulations to the alpha frequency band, and we assumed the same distribution of frequencies for all ensembles (*Ω* = 10.5 Hz; *Δ =* 1). We chose this range for three reasons. First, other MEG resting-state studies have reported significant FC in this band [7,64,68,69]. Second, earlier LSBMs have shown high correlation of simulated and empirical sFC in alpha band [17,26,47]. Third, the MEG used on this study had relatively higher power in the alpha band compared to other bands (See Supplementary Figure 6)

Other parameters were estimated with stochastic optimizers – the global coupling among ensembles, *G*, the local coupling within each ensemble, *L*_*n*_, and the spike-propagation velocity, *v*. The spike-propagation velocity and the distance between ensembles determined the interaction delays, *τ*_*ee’*_. A constant spike-propagation velocity was assumed for simplicity. However, neural fibers have a wide range of spike-propagation velocities [48,49]. Nonetheless most LSBM either neglect time delays or assume a constant velocity as well. These parameters were estimated because they have a large impact on the dynamics of LSBMs [15,16,30,47].

### Simulation scenarios and parameters estimation

The LSBM was evaluated in two scenarios. The first scenario assumed that all local couplings were identical (heterogeneous ensembles), resulting in *3* free parameters (spike-propagation velocity, global coupling, and one local coupling for all ensembles). Studies with LSBMs often assume identical ensembles, hence this scenario is similar to previous literature. In the second scenario the local couplings were free to vary across ensembles (heterogeneous ensembles), while the global parameters were fixed from the first scenario (spike-propagation velocity, and global coupling). The second scenario assumed that homotopic ensembles in the left and right hemisphere had identical local couplings, thus having *E/2* free parameters. This assumption reduced the parameter space, as well as the computing time, and it exploited the symmetries in the model. The multigraph of neural fibers and the MEG sFC are almost symmetric with respect to the interhemispheric fissure (see Figure 2). The relatively small number of parameters allowed stochastic optimizers to efficiently identify the parameters that reproduced MEG sFC. Then, trFC was computed within the restricted area of the parameter space that reproduced sFC. Supplementary Figure 1 contains an schematic representation of the parameter identification process.

#### Fitness function

The fitness function of the optimizers assessed the similarity between the sFC of MEG data and simulated neural activity. Each evaluation of the fitness function included 20 runs of the LSBM with identical parameters but different initial conditions (phases, *Ψ*_*n*_, but not *r*_*n*_). The initial phases were drawn from a uniform distribution in the range [*−π* to *π*) at the beginning of the optimization and kept across iterations of the optimizers. The initial *r*_*n*_ of each ensemble was set to its equilibrium point, 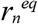, if the ensemble was detached from other ensembles 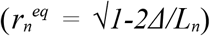. Multiple initial conditions were used to avoid overfitting of parameters to one set of initial conditions, which improves the chances of generalizing the results. Moreover, using multiple initial conditions allows for evaluating the effect of the initial conditions on the simulated FC (Supplementary Figure 5). These initial conditions might be interpreted as the previous state of the system.

The sFC from each run of the LSBM was obtained in the same way as the empirical MEG except for the orthogonalization step. Simulated data were not orthogonalized because there is no source leakage in the simulation. Another reason for not orthogonalizing was that the simulated data often were not normally distributed, which is a necessary condition for successfully applying orthogonalization [70]. Nevertheless we checked that simulated FC was not driven by zero-phase synchronization. Simulated neural activity was the imaginary part of the KOP. The same projection to neural activity has been used in prior LSBMs [15,26,27]. The median sFC across the 20 runs of the LSBM was compared with the MEG sFC by calculating the correlation between the upper triangular elements of the sFC matrices. The section *MEG and simulated data processing* provides more details on data processing.

#### Optimization constraints

The optimizers were constrained to biologically plausible solutions. These constraints prevented the LSBM from being either fully synchronized or incoherent, and enforced a minimal metastability. Full synchrony and incoherence are biologically implausible because low global synchrony appears during unconscious states [71], while high global synchrony appear during epileptic seizures [20,50]. In addition, previous studies have shown that during resting-state there is high metastability [15,21,27]. The constraints were derived from the global synchrony (Eq. 2) averaged over time, 〈 *R* 〉_*t*_, the global metastability, *SD(R)*_*t*_, the average of local synchrony over ensembles and over time, 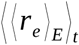, and the average local metastability over ensembles 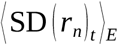. The median of these metrics across the 20 runs of the LSBM in one iteration had to comply with:

- 0.25 < median(〈 *R* 〉_*t*_) < 0.8,
- 0.05 < median(*SD(R)*_*t*_),
- 0.25 < 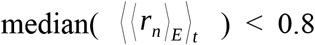,
- 0.05 < 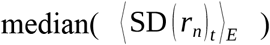

Solutions outside these constraints were heavily penalized to guarantee that the same set of parameters was not chosen again. These boundary values were selected based on visual inspection of model behavior.

#### Stochastic optimization algorithms

The optimization was carried out by three independent stochastic sampling algorithms. Several algorithms were used to examine the reproducibility and stability of solutions, and to avoid that a particular optimizer could not handle the optimization. Stochastic sampling algorithms are especially useful for complex global optimization problems in which the fitness function cannot be treated analytically. The fitness function is treated as a black box, although it is not guaranteed to find a global optimum. Nevertheless, a single best solutions was not taken, but rather a range of optimal solutions which were used to evaluate trFC later.

The optimization algorithms used were – self-adaptive Differential Evolution (aDE), Covariance Matrix Evolutionary Strategy, and Particle Swarm Optimization (PSO) [56,57,72]. These algorithms are generally successful in solving benchmark problems that include features of our problem like a high-dimensional parameter space, a non-separable function and a non-linear function [57,72–75]. Other features like the shape of the cost function could not be obtained before hand, although the chosen algorithms can cope well with unimodal and multimodal functions [57,72–75]. More information about these algorithms can be found in other sources [56,57,72,73]. The algorithms were implemented with the toolbox pagmo version 2.7 [76] in Python programming language. The 1220DE flavor of the aDE algorithm was used with all mutation variants and the iDE adaptation scheme. We opted for this algorithm because it has several parameters that are self-adapted to the features of the cost function. The parameters of the other algorithms were set to their default values, which align with the values recommended in the literature [56,57,72,73]. In the first scenario the optimizers had a population of 20 individuals and 200 generations. Each individual consisted of one set of candidate parameters for which the fitness functions was evaluated. The generations are the number of iterations of the algorithm. The optimizers on the second scenario had 110 individuals and 250 generations. The Covariance Matrix Evolutionary Strategy exhibited very poor performance, and therefore it is not reported further.

#### Model simulations

Numerical simulations of the LSBM were carried out using the time-delayed first-order Euler method. The integration step size was set to 0.001 seconds. Because of the time delays, a history of initial values was simulated independently for each ensemble from the longest time delay to the initial simulation point. Then, simulations were run for another 66 seconds in the fitness function, or 321 seconds when trFC was computed. In both cases, the first 19 seconds were discarded to remove the initial transient dynamics. The first and last second of the remaining time were discarded after filtering and Hilbert transforming the simulated neural data in order to avoid edge artifacts. In total 45 seconds were used to compute sFC in the fitness function, and 300 seconds were used for computing trFC. We kept the simulation time as short as possible to reduce the total optimization time. The model was implemented in Python. The fitness function was compiled and parallelized with Numba. One evaluation of the fitness function for one individual (i.e. 20 LSBMs, one for each of the initial conditions) took approximately 7 seconds. The 20 LSBMs in the fitness function were parallelized over 20 nodes in a computing cluster. In total one optimizer in the first scenario took approximately 8 hours. One optimizer in the second scenario took approximately 70 hours.

#### Search of time-resolved functional connectivity

trFC was searched within the range of parameters that adequately reproduced sFC. In the first scenario, trFC was evaluated within the polyhedron of parameter space defined by black lines in figure 3. A total of 1000 models were evaluated with parameters randomly taken from the optimal polyhedrons. To obtain the random parameters, first, the velocity-local-coupling polyhedron was subdivided into triangles which were uniformly sampled relative to their areas. Next, a global coupling parameter was uniformly sampled from the range of possible values given the local coupling drawn before. The polyhedrons were defined by hand to capture the set of solutions with higher sFC similarity to MEG while remaining within the biological constraints. The polyhedron approach reduced the number of trFC simulations need to cover the parameter space. Each combination of parameters was simulated 30 times with different initial phases. The initial phases were different to the ones used during the optimization. trFC as well as sFC were computed over 300 seconds of simulated data in the same way as MEG trFC and sFC. A similar approach was followed in the second scenario, apart from the polyhedron approach. In the second scenario trFC and sFC were simulated with the best 1000 parameter combinations found by each of the optimizers (2000 combinations of parameters in total). The similarity between simulated and MEG trFC was measured by the Kolmogorov-Smirnov distance between the recurrence histograms of simulated and MEG data. Similar approaches have been used before to assess the similarity of trFCs in LSBMs [26,27].

### Analysis of MEG and simulated data

The MEG resting-state datasets (300 seconds, eyes open) of 55 healthy participants, previously acquired as part of the UK MEG partnership [7,77], were used in this work. For each participant, the data were downsampled to 250Hz using an anti-aliasing filter; high-pass filtered to remove low-frequency variations below 1Hz; source-reconstructed in MNI 8mm standard space using LCMV beamforming [78] (see [77] for further details about the pre-processing). Summary time-courses were subsequently computed within each of the 68 regions of the Desikan-Killiany cortical parcellation [79] using PCA, while preserving the relative variances between regions, and signal leakage was mitigated using symmetric multivariate leakage correction [80]. To compute FC in alpha-band, the orthogonalized time-courses were band-pass filtered between 8-13Hz, their Hilbert envelope was then low-pass filtered above 0.5 Hz and downsampled to 5 Hz. Pairwise Pearson correlations between regions were computed between downsampled envelopes to obtain sFC. trFC was computed over sliding widows of 15 seconds with an overlap of 12 seconds. Within each window sFC was computed. Recurrence of sFC within-subjects was measured by the Pearson correlation between the upper diagonal elements of the sFCs for each pairs of windows. Finally, a histogram of sFC reoccurrences was built. Supplementary Figure 6 shows the time-frequency spectra of broad-band and alpha-band MEG activity.

### Computation of the anatomical network

The anatomical network was computed by averaging the results of probabilistic tractography [81,82] on 10 diffusion datasets from the Human Connectome Project [83,84] parcellated into 68 cortical regions (34 per hemisphere) using the Desikan-Killiany cortical parcellation [79]. The FSL tool ProbTrackX [85,86] was used to compute 1000 probabilistic streamlines starting from each brain voxel, and ending at the boundary between white-matter and grey-matter (WM/GM) as defined by the CIFTI format [87]. For connectivity strength between any two regions A and B was computed by dividing the number of streamlines connecting A and B, by the total number of streamlines reaching A or B (so-called fractional-scaling [88]). The resulting connectivity matrix was made symmetric by averaging with its transpose, and rescaled to have an average degree of 1. Then the connectivity matrix was normalized to have average edge weight equal to one. Finally, Euclidean distances were computed between the barycentre of each region in order to estimate the delays between them.

## Acknowledgments

- Center for Information Technology of the University of Groningen for their support and for providing access to the Peregrine high performance computing cluster
- University of Oxford Advanced Research Computing (ARC) facility http://dx.doi.org/10.5281/zenodo.22558
- Joana Cabral from University of Minho and University of Oxford for insightful discussions and instrumental role in establishing this collaboration.
- Ben Hunt from the University of Bristol (formerly in the group of Matt Brookes at the University of Nottingham) for the data collection as part of the UK MEG partnership

## Notes

### Competing Interest Statement

The authors have declared no competing interest.

