## Supplementary material for "Modulations of local synchrony over time lead to resting-state functional connectivity in a parsimonious large-scale brain model": Supplemetary Information

### Supplementary Information

#### Motivation to use the Kuramoto order parameter to describe neural synchrony.

Our LSBM is motivated by previous work which has derived exact dynamical equations for the aggregated activity of an ensemble of neurons in terms of the average firing-rate and average membrane potential using mean-field reductions from a specific set of neuron models [1–4]. A mean-field reduction allows for the simplification of the dynamics of a large number of interacting elements, in this case neurons, with a few macroscopic variables, in this case membrane potential and firing-rate. In contrast to the previous neural-mass reductions (see [5,6] for more details on neural-mass models), this mean-field reduction provides an exact bridge between the microscopic properties of individual elements and their macroscopic collective properties. For this reason, neural mean-field reductions of this sort have been referred to as 'Next generation neural mass models' [2].

Here we outline briefly the mean-field reduction of Montbrio et al. [3]. This reduction uses a quadratic integrate-and-fire (QIF) neuron model as elementary unit. It does however not rely crucially on this particular neuron type since similar reductions have been done with a theta neuron model [1,2,4]. The macroscopic membrane potential and the firing-rate of an ensemble of neurons are obtained from a heterogeneous all-to-all-connected population of QIF neurons. This reduction is exact in the thermodynamic limit, i.e., when the number of QIF neurons is infinite. The method and the steps to obtain the macroscopic variables are not within the scope of this paper, but can be found in Montbrio et al. [3]. Next we summarize this procedure.

The QIF neuron is a class I neuron model derived to simplify the Hodgkin-Huxley neuronal model. Class I neuron models go from quiescence state to periodic firing when the input current is increased, and the neuron passes through saddle-node bifurcation. The state of a QIF neuron, *j*, is given by an ordinary differential equation of its membrane potential, $V_j$.

$$\dot{V}_j = V_j^2 + I_j, \quad if \quad V_j \geq V_{th}, \quad then \quad V_r \rightarrow V_j \qquad \text{(S1)}$$

The overdot represents the temporal derivative. $I_j$ represents a trans-membrane input current. When the membrane potential reaches a threshold $V_{th}$, the neuron emits a spike and the membrane potential is reset to a resting value $V_r$. The trans-membrane input current has the form

$$I_j = \eta_j + Js(t) + I(t) \qquad \text{(S2)}$$

, where $\eta_j$ is a constant external current, $I(t)$ is a time varying input current common to all neurons, $J$ is a synaptic weight, and $s(t)$ is the mean synaptic activation.

Montbrio et. al. [3] demonstrated that the macroscopic dynamics of a fully connected ensemble of neurons in the thermodynamic limit follow exactly two ordinary differential equations in which $f$ represents the firing-rate and $v$ the average membrane potential.

$$\dot{f} = \gamma / \pi + 2 f v \tag{S3}$$
$$\dot{v} = v^2 + \overline{\eta} + Jf + I(t) - (\pi f)^2 \tag{S4}$$

The heterogeneity of the external currents is represented by a Lorentzian distribution with half-width at half-maximum $\gamma$ and maximum peak at $\eta$.

Furthermore, Montbrio et al. [3] demonstrated that the macroscopic dynamics of the ensemble can be simplified further with a conformal mapping to the Kuramoto order parameter (KOP), $z$. Note that other mean-field reductions with the $\theta$-neuron model can be mapped to the KOP as well [1,2]. This mapping has the form of

$$z = \frac{1-w}{1+w}, \quad w = \frac{1-z}{1+z}, \quad \text{where} \quad w \equiv f\pi + iv, \quad \text{and} \quad z = r\, e^{i\psi} \quad \text{with} \quad r = [0,1], \quad \text{and} \quad \psi = [-\pi, \pi]. \tag{S5}$$

In the above equation, the asterisk denotes the complex conjugate. In our LSBM we use the KOP as the descriptor of the neural activity in an ensemble of neurons, although the dynamics of the KOP are not given by the equations of Montbrio et al. [3].

#### Derivation of the LSBM equations

For the sake of simplicity we obtain the KOP from a fully connected network of Kuramoto oscillators [7,8] instead of equations 3S, 4S, and 5S. The Kuramoto model has been widely studied theoretically and it has also been applied to many biological examples of collective synchronization [8]. In a Kuramoto model the synchronization is controlled by a coupling parameter. Natural frequencies are drawn form a probability distribution $g(\omega)$ and initial phases are drawn from a uniform distribution ($r = 0$). When the coupling is low, each oscillator follows its own natural frequency. When the coupling increases above a critical value the oscillators began to synchronize. As the coupling continues increasing their synchrony increases until they reach full synchrony if the coupling continues increasing.

Next, we derive the equations of the LSBM. We consider a network with $E$ ensembles labeled $n = 1,2, \ldots E$, where ensemble $n$ contains $N_n$ oscillators $\theta_n^i(t)$ with $i = 1,2, \ldots N_n$. Pairs of ensembles, $n$ and $p$, are coupled through the anatomical network $A_{np}$ (the connectome), likewise, $\tau_{np}$ represents the time delays between the same pair of ensembles. The coupling between all pairs of ensembles is scaled by a global coupling $G$. The global coupling is scaled by 1/E, in order to impose community structure with stronger local than global coupling. Local couplings, $L_n$, determine the coupling within each individual ensemble of oscillators. The local couplings allow us to manipulate the synchrony within each ensemble. For simplicity, the anatomical network $A$, the global coupling scaling factor $G/E$, and the local couplings $L_n$ are embedded into an $E \times E$ coupling matrix $K_{\mathrm{np}}$. The diagonal elements of the matrix $K_{np}$ contain the local couplings $L_n$. The off-diagonal elements of $K_{\mathrm{np}}$ contain the values of the anatomical network $A$ scaled by the global coupling scaling factor $G/E$. The natural frequencies of the oscillators in each ensemble are drawn from different probability distributions. The distribution of natural frequencies in one ensemble is denoted by $g_n(\omega)$, and the natural frequency of one oscillator in the same ensemble is $\omega_n^i$. With the previous definitions, the evolution of oscillator $i$ in the ensemble $n$ follows the next delayed differential equation.

$$\dot{\theta}_n^i = \omega_n^i + \sum_{p=1}^{E} \frac{K_{np}}{N_p} \sum_{j=1}^{N_p} \sin\left(\theta_p^j\left(t-\tau_{np}\right) - \theta_n^i(t)\right) \tag{S6}$$

We are interested in describing each ensemble by the KOP introduced in equation S5. The KOP in an ensemble of oscillators can be defined as follows [7]:

$$z_n = r_n e^{i\psi_n} = \frac{1}{N_n} \sum_{i=1}^{N_n} e^{i\theta_n^i}, \tag{S7}$$

where $r_n$ and $\psi_n$ represent the average phase coherence and the mean phase in ensemble $n$ respectively. Substituting equation S7 into equation S6, we can rewrite equation S6 in terms of the KOP as follows:

$$\theta_n^i = \omega_n^i + \sum_{p=1}^{E} K_{np} \operatorname{Im}\left(e^{-i\theta_n^i(t)} z_p\left(t-\tau_{np}\right)\right) \tag{S8}$$

Now we consider the case in which the number of oscillators in one ensemble is very large i.e. $N_n \to \infty$, and we define $\rho_n(\theta_n, \omega_n, t)$ as the probability density of oscillators in $n$ with phase $\theta$ and frequency $\omega$ at time $t$. Since the number of oscillators on each ensemble remains unchanged, $\boldsymbol{\rho_n}$ satisfies the local continuity equation,

$$\frac{\partial \rho_n}{\partial t} + \frac{\partial}{\partial \theta_n}\left(\rho_n, \dot{\theta}_n\right) = \frac{\partial \rho_n}{\partial t} \frac{\partial}{\partial \theta_n}\left\{\rho_n\left[\omega_n + \sum_{p=1}^{E} K_{np} \operatorname{Im}\left(e^{-i\theta_n(t)} z_p\left(t-\tau_{np}\right)\right)\right]\right\} = 0 \tag{S9}$$

Next we consider the Fourier series of $\rho_n$

$$\rho_n(\theta,\omega,t) = \frac{g_n(\omega)}{2\pi}\left[1+\sum_{k=1}^{\infty} c_{n,k}(\omega,t)e^{ik\theta} + c.c.\right], \tag{S10}$$

where c.c. stands for complex conjugate. Following the Ott and Antonsen ansatzt [9], we consider a class of $c(\omega,t)$ such that $c_{n,k}(\omega,t) = a_n(\omega,t)^k$, where $|a_n(\omega,t)| \leq 1$. Then, inserting the series expansion $a_n(\omega,t)$ into equation S9 we obtain

$$\frac{\partial a_n}{\partial t} + i\omega_n a_n + \frac{1}{2}\sum_{p=1}^{E} K_{np}\left[z_p(t-\tau_{np})a_n^2 - z_p(t-\tau_{np})\right] = 0. \tag{S11}$$

Letting the distribution of frequencies $g_n(\omega)$ to be a Lorentzian distribution with center at $\Omega_n$ and spread of $\Delta_n$, i.e.

$$g_n(\omega) = \frac{\Delta_n}{\pi\left[(\omega - \Omega_n)^2 + \Delta_n^2\right]} \tag{S12}$$

we can calculate $z_n$ in the continuum limit as

$$z_n = \int_{-\infty}^{\infty}\int_0^{2\pi} \rho_n(\theta,\omega,t)e^{i\theta}d\theta d\omega = \int_{-\infty}^{\infty} g_n(\omega)a_n(\omega,t)d\omega = a_n(\Omega_n - i\Delta_n,\mathrm{t}) \tag{S13}$$

Thus, by evaluating equation S11 at $\omega = \Omega_n - \Delta_n$, we can obtain the dynamics of $z_n$ as

$$\dot{z}_n + (\Delta_n - i\Omega_n)z_n + \frac{1}{2}\sum_{p=1}^{E} K_{np}\left[z_p(t-\tau_{np})z_n^2 - z_p(t-\tau_{np})\right] = 0 \tag{S14}$$

Therefore the LSBM is defined by $E$ complex delayed differential equations. Because of the first equality of equation S7, the LSBM with $E$ equations can be converted to $2E$ real delayed differential equations with the forms

$$\dot{r}_n = -\Delta_n r_n + \frac{1-r_n^2}{2}\sum_{p=1}^{E} K_{np}\mathrm{Re}\left[z_p(t-\tau_{np})e^{-i\psi_n}\right] \tag{S15a}$$

$$\dot{\psi}_n = \Omega_n + \frac{r_n^2+1}{2r_n}\sum_{p=1}^{E} K_{np}\mathrm{Im}\left[z_p(t-\tau_{np})e^{-i\psi_n}\right] \tag{S15b}$$

where $\dot{r}_n$ and $\dot{\psi}_n$ describe the temporal evolution of the KOP ( $z_n = r_n e^{i\psi_n}$ ) in terms of the level of synchrony and the mean phase in ensemble $n$ respectively. The equations 1a and 1b in the main text are obtained by extracting the global coupling scaling factor, $G/E$, the local coupling, $L_n$, and the anatomical network of neural fibers, $A$, from the coupling matrix, $K_{np}$ , in equations S15a and S15b.

#### Supplementary figures

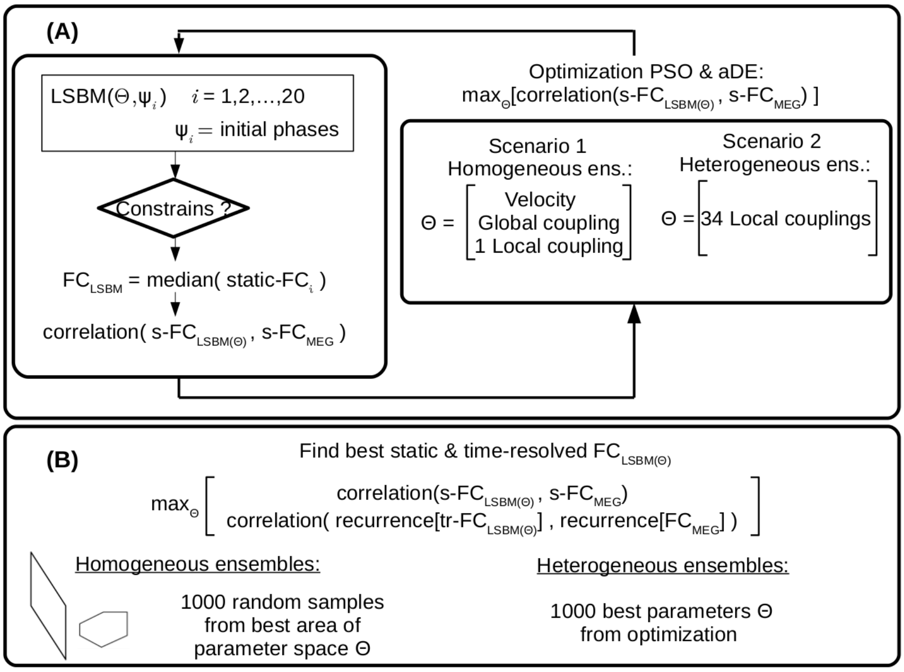


**Supplementary Figure 1. Schematic representation of the parameter identification process.** (A) Optimization of parameters for the first (homogeneous ensembles) or the second scenario (heterogeneous ensembles). (B) Simulations to find the parameters that are able to predict static FC and time-resolved FC.

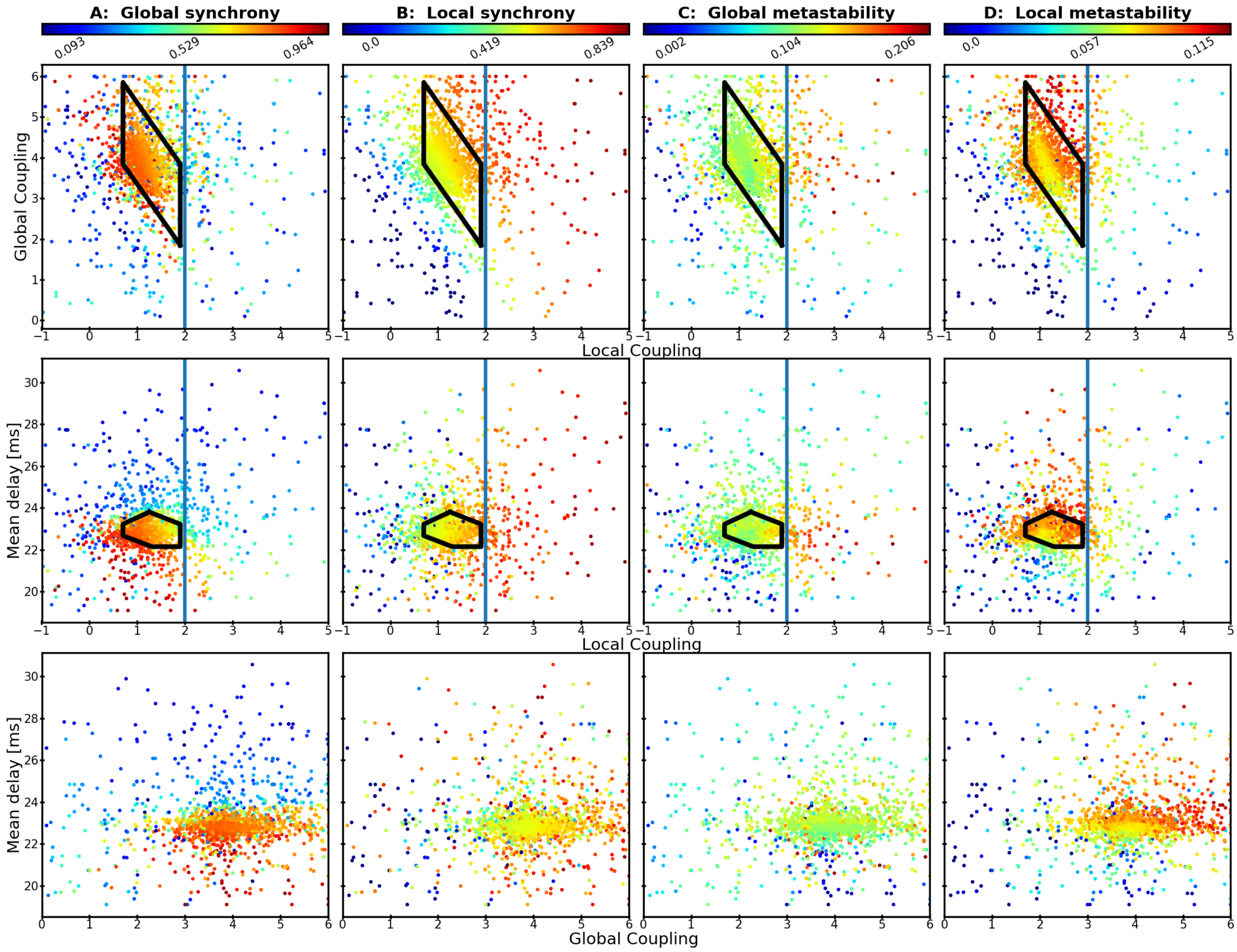


**Supplementary Figure 2. Dynamics of the model with homogeneous ensembles.** Each dot corresponds to one combination of parameters (x-y axis) during the optimization. Each column corresponds to a different dynamical feature. (A) Global synchrony averaged over time. (B) Local synchrony averaged over ensembles first and then over time. (C) Global metastability (D) Local metastability averaged over ensembles. For each metric, each value corresponds to the median obtained across 20 simulations computed with different initial conditions. Vertical blue lines indicate the critical local couplings of the ensembles, $L^c$.

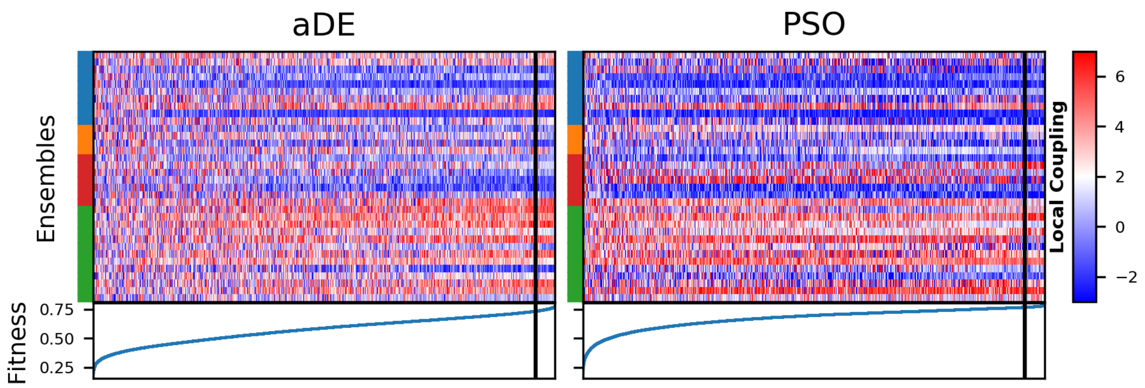


**Supplementary Figure 3. Evolution of local couplings optimization sorted by fitness to MEG sFC.** Each row inside the upper panel depicts the local couplings of one ensemble. The columns correspond to iterations of the respective optimizer (aDE and PSO). The brain lobe associated with each ensemble is color coded on the left (same as Figure 2). The fitness (Pearson correlation between MEG and simulated sFC) of each iteration is at the bottom of the panels. trFC was evaluated at the parameters to the right of the black vertical line.

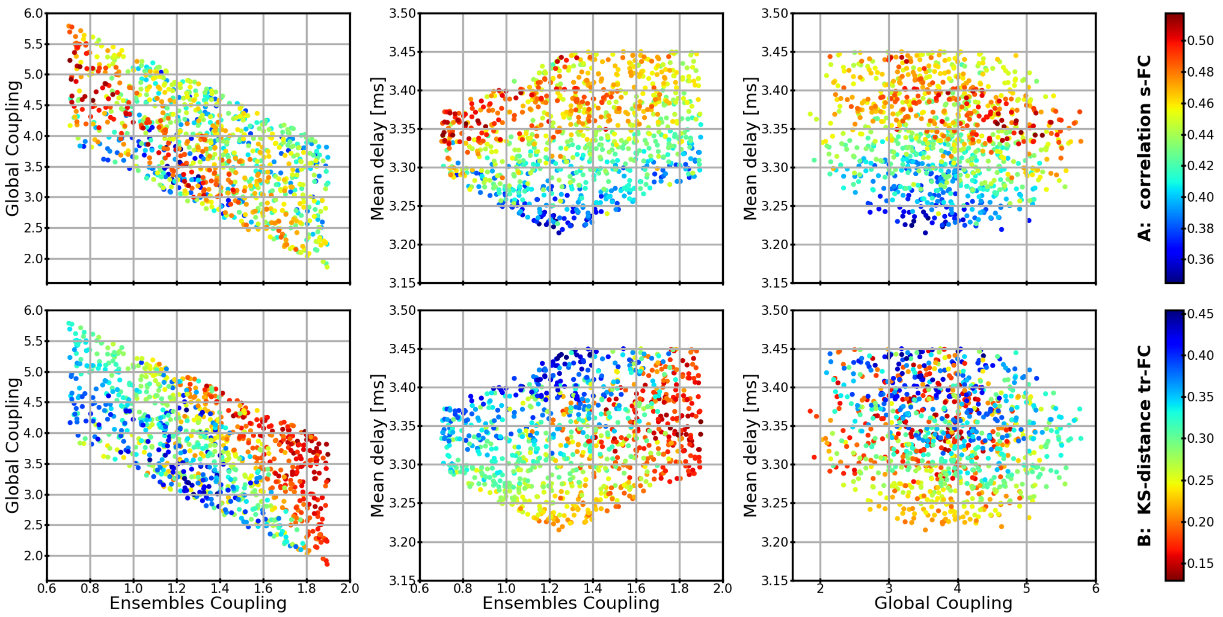


**Supplementary Figure 4. FC comparison between 300-second simulations and MEG data.** Each dot corresponds to one combination of parameters – global coupling, local coupling, and mean delay. The parameters are restricted to the area that gave the best sFC predictions during the optimization (black area in figures 2 and 3). (sFC, first row) correlation between the simulated and MEG sFC. (trFC, second row) KS-distance between the histograms (simulated vs. MEG data) built from the recurrence of sFC recurrence over a 15-second sliding window with 12-second overlap. Each dot is median of 30 simulations with identical parameters and different initial conditions.

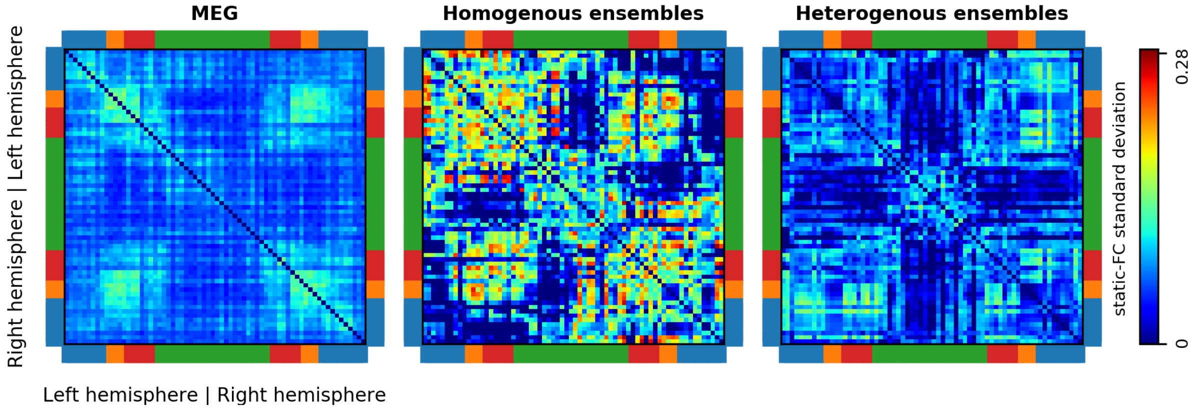


**Supplementary Figure 5. Functional connectivity (FC) variability.** (Left) Standard deviation of the FC over subjects (Middle, model with homogeneous ensembles) Standard deviation of the FC from simulations with identical parameters but different initial conditions. (Right, model with heterogeneous ensembles) Standard deviation of the FC from simulations with identical parameters but different initial conditions.

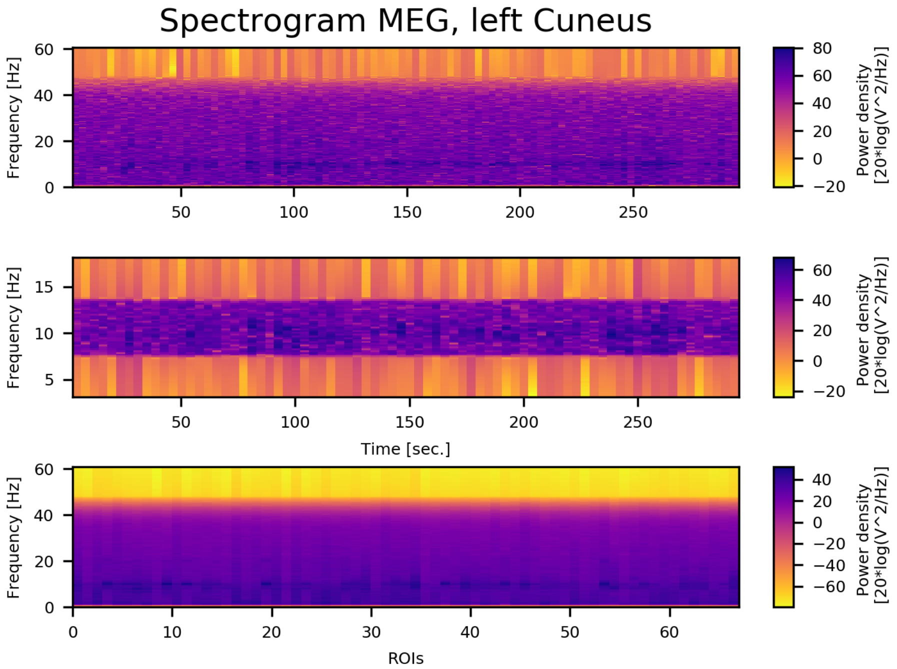


**Supplementary Figure 6. time-frequency spectrogram from MEG activity in the alpha band.** (*Above*) Broadband activity with a high-pass filter at 48 Hz. (*Middle*) Band-pass filtered alpha-band activity. (*Bottom*) Welch periodogram of broadband activity in all ROIs .

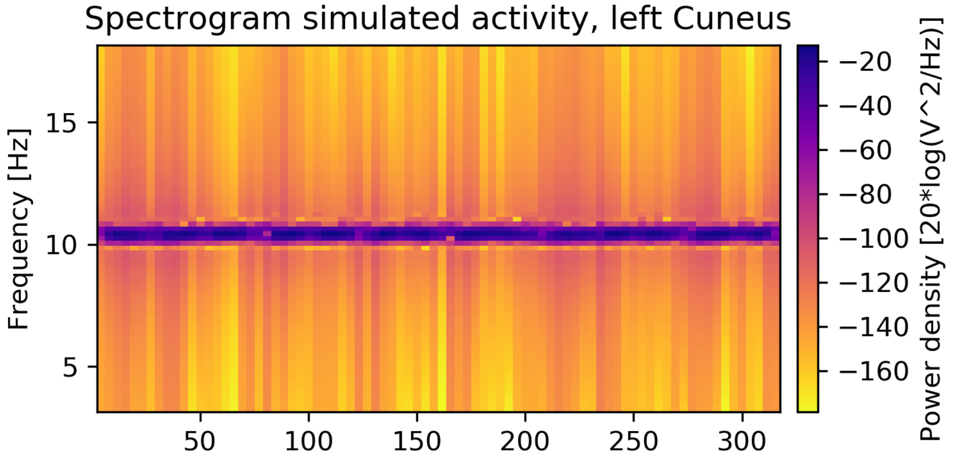


**Supplementary Figure 7. time-frequency spectrogram from simulated activity in the alpha band.** The parameters are the same parameters as Figure 2 and 5 for heterogeneous ensembles.
